# Sperm attraction by female reproductive fluid in a fish with an unconventional fertilisation strategy

**DOI:** 10.64898/2026.02.27.708519

**Authors:** Alexandra Glavaschi, Matej Polačik, Martin Reichard

## Abstract

Active sperm guidance towards the eggs is necessary for gametes to meet within the fertilisation window and is usually achieved through the action of female reproductive fluids (FRF hereafter) on sperm flagellar movement. However, it remains unknown whether FRF retains this function in species with unusual fertilisation strategies. Such examples include systems where gametes are released within confined microhabitats characterised by flow-driven physical processes that could facilitate sperm-egg encounters, hence selection on active sperm guidance may be relaxed. Alternatively, sperm attraction could be maintained by pleiotropic effects of genes involved in other reproductive functions of FRF. Here we tested whether FRF of species with unusual modes of fertilisation retain sperm attractant properties using the bitterling, a small cyprinid fish that parasitises freshwater mussels. Bitterling deposit sperm and eggs inside the mussel gills and gamete interactions are facilitated by the water current generated by mussel respiration. Using a recently developed sperm selection chamber and the European bitterling, we find that more sperm accumulate in the FRF channel compared to the water control. Moreover, European bitterling sperm showed no preference for conspecific FRF over that of a distantly related Asian bitterling FRF. Our results suggest that bitterling FRF resembles that of species with conventional fertilisation modes, implying that it could mediate sperm selection and cryptic female choice. We discuss alternative evolutionary scenarios underlying the persistence of sperm attractant properties of bitterling FRF despite the shift to a physically mediated fertilisation environment.

## Introduction

Spermatozoa guidance mechanisms are crucial to ensure sperm-egg encounters within the fertilisation window in often hostile environments (Kholodnyy et al., 2020). Sperm attraction towards the eggs is usually achieved through the influence of active compounds contained in female reproductive fluids, such as amino-acids, peptides and proteins, on sperm flagellar movement (Kholodnyy et al., 2020, Yoshida et al., 2008).

Female reproductive fluids (FRF hereafter) are media derived from the female reproductive tract, accessory glands or eggs with which sperm interacts before and during fertilisation (Gasparini et al., 2020a). In internal fertilisers, FRFs are kept within the female tract while in external fertilisers they are released together with the eggs. The primary functions of FRFs are to protect and nourish the maturing oocytes and eggs and to provide the optimal conditions for fertilisation (Aguilar and Reyley, 2005). Proteomic analyses in various taxa indicate that FRFs guard against oxidative stress (Baer et al., 2009) and pathogens (Johnson et al., 2014). Additionally, in external fertilisers FRFs buffer against adverse environmental factors and lengthen egg fertilisation window (Pinzoni et al., 2023, Dietrich et al., 2012, Gueho et al., 2024). Irrespective of fertilisation mode and origin (ovarian, coelomic, follicular etc.), FRFs are the media through which sperm must swim to reach and fertilise the eggs, and a growing body of evidence shows that FRFs are not passive environments with respect to sperm function, but influence a whole suite of sperm traits (Cattelan et al., 2023, Hadlow et al., 2023, Poli et al., 2019) with consequences for fertilisation success (Pinzoni et al., 2024).

The sperm attractant role of FRFs has been identified in a wide variety of internal and external fertilisers (Lymbery et al., 2017, Serrano et al., 2001, Fitzpatrick et al., 2020, Kholodnyy et al., 2020), yet it remains unknown if this function extends to species with unconventional modes of fertilisation. Such examples concern reproductive parasites or species where fertilisation is aided by the environment, but close interactions between gametes and surrounding fluids remain unexplored in these taxa.

Our main aim in this study is to investigate the sperm attractant role of FRF in bitterling, a group of freshwater fish with a long (c. 32 million years) evolutionary history as reproductive parasites of mussels (Zhao et al., 2016). Bitterling are small cyprinids distributed across East Asia and Europe. During spawning season, females develop an ovipositor – a long tubular flaccid organ which they use to deposit eggs in the mussel gill cavity. This is achieved by pushing the base of the ovipositor inside the mussel exhalant syphon, at which point internal pressure forces the eggs, together with a small volume of fluid, through the length of the ovipositor until they are released into the water tubes of the mussel gill. Males ejaculate over the inhalant syphon and the water current carries the sperm through the gill membrane, where eggs are fertilised and embryos develop for approximately one month (Smith et al., 2004). Thus, bitterling are external fertilisers in the sense that fertilisation takes place outside of the female body, yet sperm and eggs are not released into the open water but contained within the mussel cavity and gamete encounters are facilitated by mussel respiration. Bitterling are completely dependent on mussels for reproduction, such that visual and olfactory cues of mussel presence are essential for both males and females to get into spawning condition (Smith et al., 2004, Heschl, 1989). Males commonly ejaculate over the mussel before oviposition has taken place, usually repeatedly (Smith et al., 2014), so eggs are often deposited in an environment already containing activated sperm. Whether or not bitterling FRF retains sperm attractant properties is not straightforward to predict. On the one hand, because gamete interactions are mediated by the physical processes involved in mussel respiration, we would argue that “ancestral” sperm attractant properties of bitterling FRF are redundant and should be lost in modern species. However, if the same active compounds are responsible for the effects of FRF on both eggs and sperm, then sperm attraction could be maintained as a by-product of the primary roles of FRF.

In this study, we use the European bitterling *Rhodeus amarus* and a recently developed multi-channel sperm selection chamber (Devigili et al., 2021) to test the sperm attractant properties of bitterling FRF. If sperm attractant properties of bitterling FRF are conserved, we predicted that it is so because FRF retains influence on sperm function more generally. To test this prediction, we investigated the boosting effect of FRF on sperm longevity, another attribute of FRF across numerous taxa (Zadmajid et al., 2019, Hadlow et al., 2023) that could potentially be lost in bitterling. This is because the number of pre- rather than post-oviposition ejaculations is associated with fertilisation success in natural spawnings of *R.amarus* (Reichard et al., 2004) and bitterling sperm is unusually long-lived compared to that of other external fertilisers (Browne et al., 2015). We therefore measured *R. amarus* sperm motility duration in a FRF solution and water control.

Across taxa, the effects of FRFs on sperm function are species-specific (Myers et al., 2020, Yeates et al., 2013) and, as mediators of cryptic female choice, FRFs contribute towards the maintenance of reproductive barriers (Kustra et al., 2025). The bitterling radiation contains at least 60 species (Chang et al., 2014), often coexisting across China, Korea and Japan and using the same mussel species as hosts for their eggs (Kitamura, 2007), with the European bitterling being the only lineage in the West Palearctic. All bitterling species share the same fertilisation strategy, meaning that sperm and eggs are deposited inside a mussel gill cavity. If the sperm attraction property is retained by bitterling FRF, one possible reason behind it is their role in species-specific sperm recognition. Our second aim was to test whether the sperm attractant properties of bitterling FRF, if any, are species-specific or general. With the same sperm selection device, we used sperm of *R. amarus* and FRF from a distantly related allopatric species, *Sinorhodeus microlepis,* to compare sperm behaviour between (a) heterospecific FRF and water and (b) conspecific and heterospecific FRF.

## Materials and methods

### Study animals

European bitterling *R. amarus* originated from two populations: Kyjovka river in the southeast of the Czech Republic (48°47’N, 17°01’E) and Ballica river in Turkey (41°00’N, 29°25’E). *S. microlepis* were F1 descendants of 11 adult fish imported from the Yangtse River basin, Chongqing City, China (29°33′N, 106°33′E).

During the breeding season (April-June for *R. amarus* and April-September for *S. microlepis*), fish are housed in 1000L outdoor ponds with access to mussels and natural food, and additionally fed a mixed diet of flake food, pellets and bloodworms six days a week. For overwintering, both species are kept in a designated facility with a glass ceiling, naturally short daytime and temperatures ranging from 6 to 15°C.

### FRF and sperm solution preparation

Females in spawning condition, identified by the presence of a fully developed ovipositor, were anesthetised in clove oil and placed on a dry glass petri dish, the ovipositor fully extended and dried using a paper tissue and the eggs and FRF released by applying gentle pressure to the abdomen. A volume of distilled water equivalent to 3μl/egg was added to the clutch, the eggs carefully rinsed and the resulting diluted FRF transferred to an Eppendorf tube using an insulin syringe.

Males were anesthetised in clove oil, the vent areas dried with paper tissue and the ejaculate released by applying gentle pressure to the abdomen, collected with a pipette and transferred to an Eppendorf tube. To reduce bias arising from male-by-female interactions (Daupagne et al., 2024), we diluted 1μl ejaculate from each of two males into 1ml NaCl 0.9% for a concentration of 2μl/1ml. Adjusting sperm concentration to a standardised value prior to the assays was logistically challenging, therefore we counted the number of sperm from 3μl solution and included this value as a covariate in all analyses (see below). Sperm and FRF were kept on ice and used within five hours of collection. Pilot tests indicated that sperm remain viable in this time window.

### Sperm attraction assays

Sperm attraction procedures closely follow the protocol described in (Devigili et al., 2021). The sperm selection chambers used for assays are small rectangular devices featuring three small wells connected through channels to a central well (Figure 1). Detailed descriptions of the chambers, including all dimensions and the files used for 3D printing, are available as supplementary materials in (Devigili et al., 2021). We 3D printed three chambers made of biologically inert resin (Liqcreate Bio-Med Clear, https://www.liqcreate.com/product/bio-med-clear-biocompatible-resin/) using the lost wax technique (Devigili et al., 2021).

**Figure 1.**
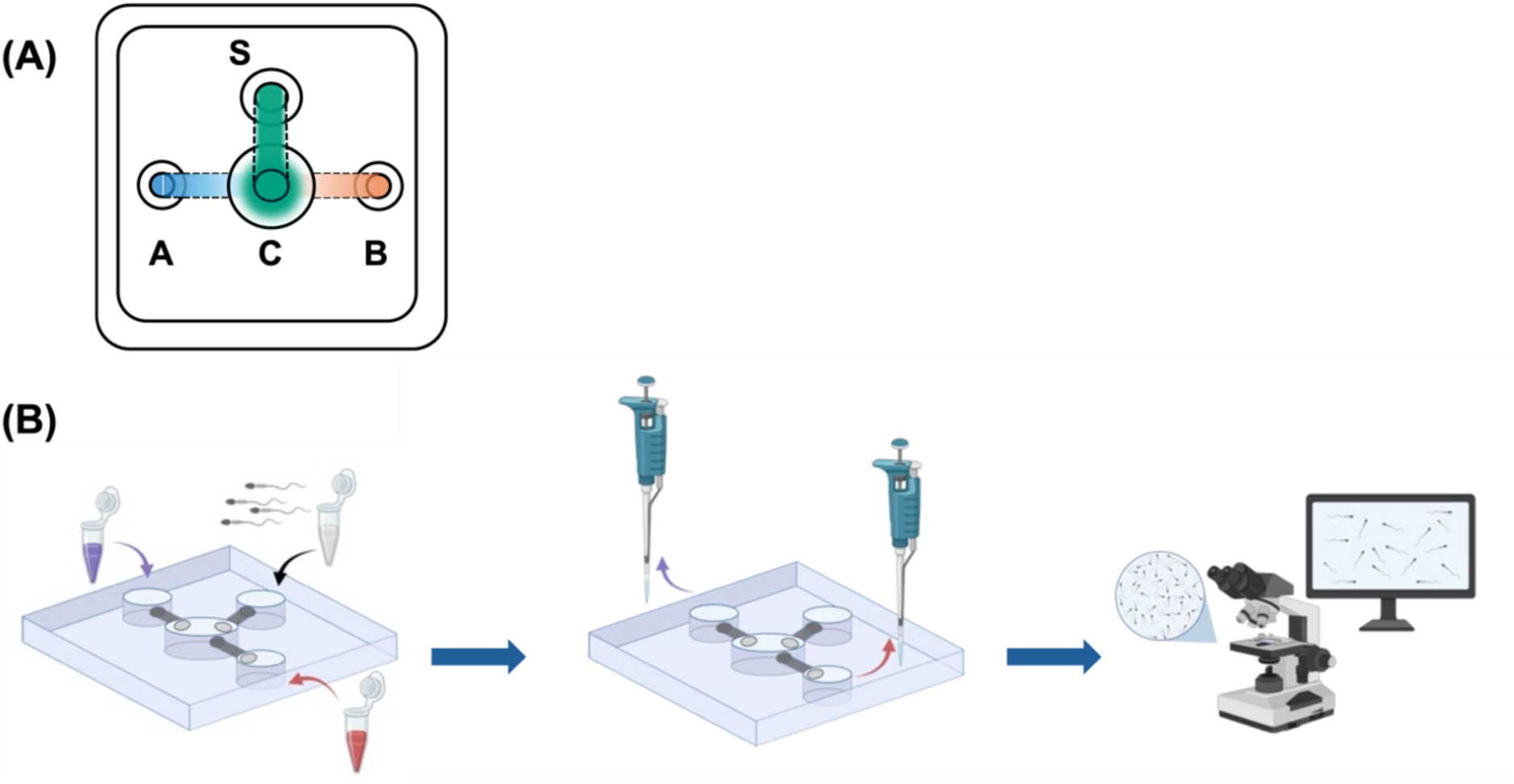
**(A)** Schematic representation of the sperm selection chamber (not to scale). A and B: the wells where the media (FRF and water) are loaded. S: well where the sperm solution is loaded. C: central well where sperm encounters the FRF-water (or conspecific-heterospecific) gradient. **(B)** Summary of the experimental procedure. First, the two media are added into wells A and B and allowed to form a gradient in well C, after which the sperm solution is added in well S. After a predetermined amount of time, 3μl liquid are collected from wells A and B in the same order as loading and the number of sperm in each sample is counted. Created with Biorender.com.

We conducted three independent types of assays to test *R. amarus* sperm attraction to: 1) conspecific FRF *vs* water, 2) heterospecific FRF *vs* water, 3) conspecific FRF *vs* heterospecific FRF. Replication units consist of unique FRF – sperm pool combinations and the sample breakdown is shown in Table 1. We ran two trials for each FRF – sperm pool combination, with the order of media loading and sperm retrieving and counting randomly predetermined and balanced.

**Table 1.** Sample size breakdown for each type of sperm attraction trial. Due to limited availability, certain individuals were used in multiple sperm pools, pairs or triads, leading to apparently unbalanced sample size for the number of fish and number of trials.

| Trial type | *R. amarus*<br>females | *S. microlepis*<br>females | *R. amarus*<br>males | *R. amarus*<br>male pairs | Total number<br>of trials |
| --- | --- | --- | --- | --- | --- |
| Conspecific FRF | 14 | NA | 20 | 11 | 21 x 2 |
| Heterospecific FRF | NA | 15 | 19 | 9 | 17 x 2 |
| Con- vs<br>Heterospecific FRF | 8 | 8 | 12 | 8 | 12 x 2 |

Trials consisted of the following steps, summarised in Figure 1B. First, to avoid the formation of air bubbles, 12μl distilled water were loaded into each of the three small wells (S, A and B; Figure 1A). Then, 5μl of the medium of interest (water, *R. amarus* FRF or *S. microlepis* FRF) were added into wells A and B, respectively. The liquids were allowed to form a gradient in the central well for 90 seconds, after which 20μl sperm solution were loaded into well S. Upon activation in water, sperm starts swimming towards the central well where it encounters the FRF-water or conspecific-heterospecific FRF gradient. After 50 seconds from sperm loading, 3μl liquid were collected from wells A and B and transferred to Eppendorf tubes. Sperm cells were then counted from each sample on a Makler chamber under an Olympus BX53 phase contrast microscope. Sperm counting from the media retrieved from wells A and B was performed in the same order as loading and collecting. Sperm selection chambers were cleaned between trials by flushing with distilled water and drying with compressed air.

### Sperm swimming longevity measurement

We used *R. amarus* to measure sperm longevity in a FRF solution prepared as above and a water control. An insulin syringe needle was gently dipped into the undiluted ejaculate then rinsed in 3μl medium previously loaded onto a Makler chamber under a phase contrast microscope. Longevity was measured as time elapsed from activation until sperm cells stopped swimming entirely. We ran 13 repeated trials using unique male-female pairs, with the order between FRF and water pre-determined and balanced.

### Data analysis

We investigated whether the number of sperm retrieved from the focal well (*R. amarus* or *S. microlepis* FRF) was different from the control well (water or *S. microlepis* FRF, respectively). We ran three separate general linear mixed-effects models with Gaussian distribution and identity link function using the *glmmTMB* package (McGillycuddy et al., 2025). The number of sperm cells retrieved from the two wells was the response variable in all models. We applied a square root transformation for the conspecific FRF *vs* water contrast and an arcsine transformation to the con- *vs* heterospecific FRF contrast to achieve a Gaussian distribution of the response variable. All models included medium (FRF or water) as a fixed factor, female-sperm pool combination as a random factor to account for the paired design, and sperm concentration as a covariate.

We used the same model structure as above to compare sperm swimming longevity in FRF and a water control. Sperm longevity was log-transformed to meet model assumptions. The medium (FRF or water) was included as a fixed factor and the male-female pair ID as a random factor. We calculated marginal and conditional *R*^2^ to compare the variance explained by fixed and random effects (Nakagawa et al., 2013).

We used the *DHARMa* diagnostics package (Hartig, 2016) to check model fits and we further examined overdispersion using the *check_overdispersion* function of the *performance* package (Lüdecke et al., 2021). We carried out all analyses in R (version 4.5.0) and we report means and standard errors (s.e.) throughout.

## Results

Bitterling sperm clearly responded to the chemoattractant properties of conspecific FRF. When tested against a water control, we retrieved a higher number of sperm cells from the FRF well in 19 out of 21 trials (Figure 2A, B), and this difference was statistically significant (FRF: 143.09±10.48 water: 77.64±10.22 ; Table 2). Bitterling sperm was also attracted towards heterospecific FRF. The wells loaded with *S. microlepis* FRF attracted significantly more *R. amarus* sperm compared to the water control (FRF: 137.11±10.6; water: 105.94±11.55; Table 2; Figure 2C, D).

**Figure 2.**
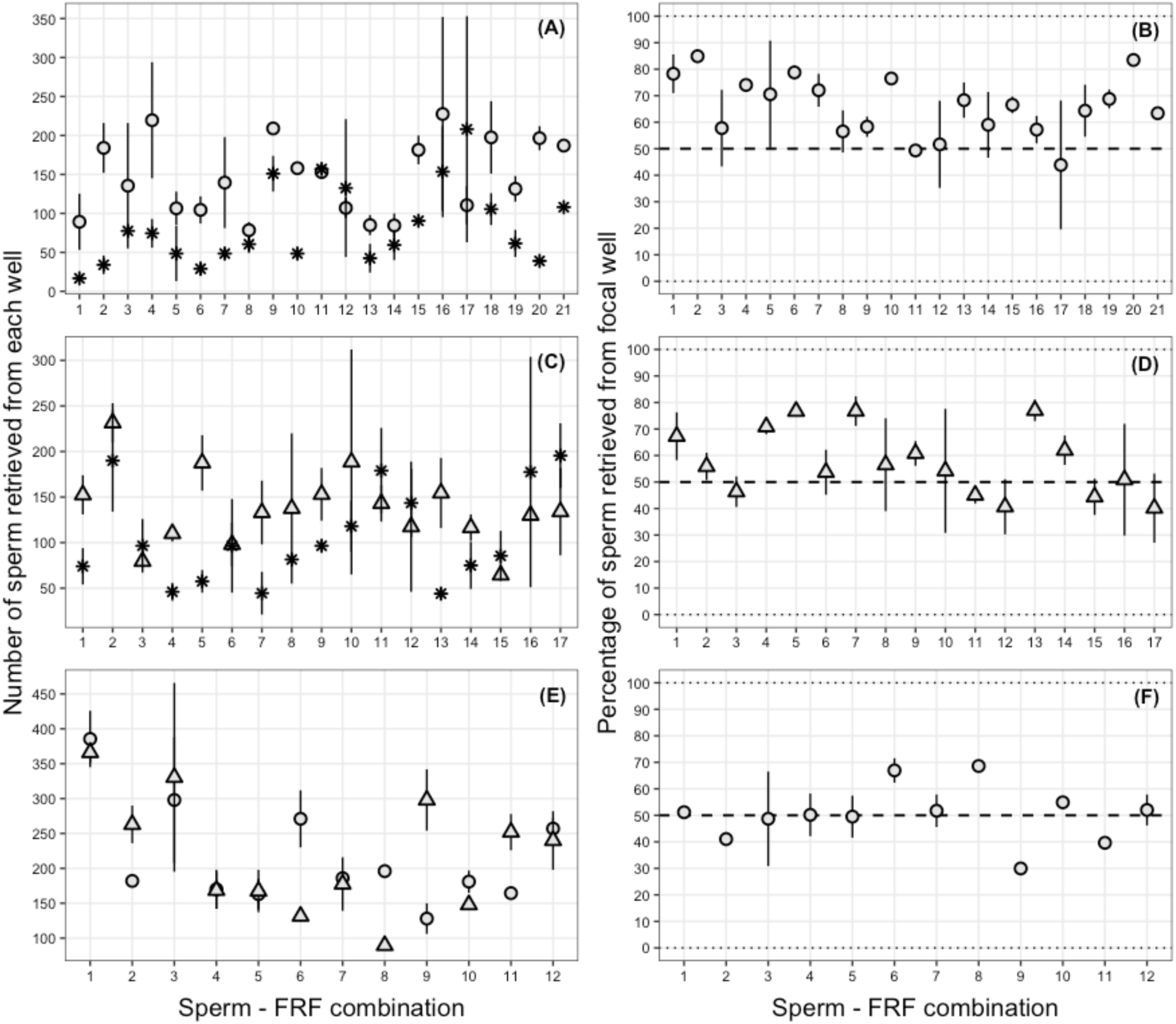
Mean (+/- s.e.) number of sperm retrieved from the focal and control well for each individual sperm pool-FRF combination (A, C, E). Percentage of sperm (+/- s.e) collected from the focal well out of the total number of sperm cells collected from both wells (B, D, F). Water is represented as asterisks, *R. amarus* FRF as circles and *S. microlepis* FRF as triangles.

**Table 2.** Results of general linear mixed-effects models testing for differences in the number of sperm cells collected from the focal and control wells of the selection chambers.

|  | Estimate | Std error | z | p |
| --- | --- | --- | --- | --- |
| (1) Conspecific FRF vs water |  |  |  |  |
| Intercept | 10.174 | 1.09 | 9.271 | <0.001 |
| Medium | -3.44 | 0.561 | -6.126 | <0.001 |
| Sperm concentration | 0.005 | 0.003 | 1.651 | 0.09 |
| $R^2_{\text{conditional}} = 0.439$; $R^2_{\text{marginal}} = 0.290$ | | | | |
| Random effects | $\sigma^2_{(\text{Combination})} = 1.762$ | | | |
| (2) Heterospecific FRF vs water |  |  |  |  |
| Intercept | 130.916 | 26.414 | 4.956 | <0.001 |
| Medium | -34.75 | 15.881 | -2.188 | 0.028 |
| Sperm concentration | 0.015 | 0.049 | 0.311 | 0.756 |
| $R^2_{\text{conditional}} = 0.072$; $R^2_{\text{marginal}} = 0.072$ | | | | |
| Random effects | $\sigma^2_{(\text{Combination})} = 0$ | | | |
| (3) Con- vs heterospecific FRF |  |  |  |  |
| Intercept | 5.8 | 0.379 | 15.295 | <0.001 |
| Medium | -0.015 | 0.088 | -0.178 | 0.859 |
| Sperm concentration | 0.00 | 0.001 | 0.563 | 0.573 |
| $R^2_{\text{conditional}} = 0.349$; $R^2_{\text{marginal}} = 0.014$ | | | | |
| Random effects | $\sigma^2_{(\text{Combination})} = 0.048$ | | | |

Bitterling sperm attraction to FRF was not species-specific. When *R. amarus* FRF was compared to *S. microlepis* FRF, we found no difference in the number of cells collected from the two wells (*R. amarus* FRF: 215±16.25; *S. microlepis* FRF: 219.33±19.7; Table 2; Figure 2E, F). Sperm concentration had no effect on the number of cells retrieved in any of the trials, nor did the identity of the female-sperm pool combination (Table 2). Random effects (individual identities of males and females) explained substantial proportions of variation in the conspecific and con- vs heterospecific trials (Table 2).

Sperm swimming longevity in *R. amarus* was enhanced by FRF. The average motility duration in a FRF solution was more than double compared to a water control (FRF: 243.38±9.25 s; water: 102.69±10.74 s; General Linear Model: estimate (±s.e.) = -0.911±0.089; z=-10.13; p<0.001; Figure 3).

**Figure 3.**
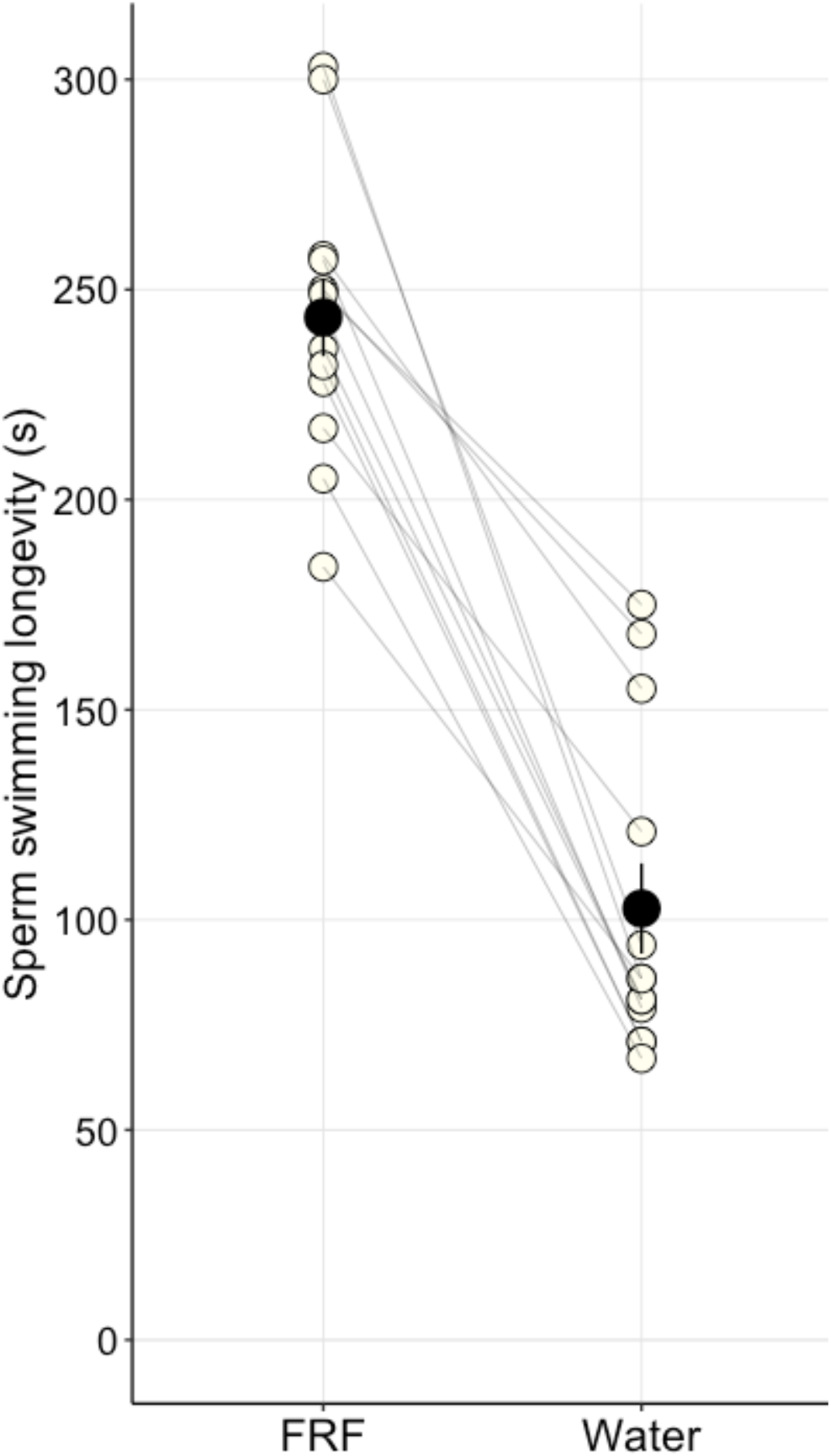
*R. amarus* sperm longevity duration in a FRF solution compared to a water control. Open circles connected by lines are measurements for individual male-female pairs and filled circles are averages per treatment (± s.e.).

## Discussion

Despite the European bitterling being a model system for sperm competition (Candolin and Reynolds, 2002, Smith et al., 2003, Smith et al., 2014), fine-scale fertilisation dynamics in this taxon remain poorly studied. This is a significant knowledge gap, because understanding gamete interactions in bitterling in relation to the mussel environment could help to understand the evolution of novel fertilisation strategies.

We found that significantly more European bitterling sperm accumulates in the FRF channel of the sperm selection chamber than in the channel containing a control solution. This includes the FRF of an unrelated bitterling species, although the response was weaker compared to that of conspecific FRF (Figure 2). This result confirms that the sperm attractant property of FRFs (Lymbery et al., 2017, Devigili et al., 2021, Evans and Sherman, 2013, Yeates et al., 2013, Yoshida et al., 2002, Fitzpatrick et al., 2020, Serrano et al., 2001) is not lost in the bitterling, a species where gamete encounters are primarily mediated by physical, flow-driven processes.

FRFs have been the focus of much recent research due to their roles as mediators of post-mating sexual selection (Alonzo et al., 2016, Gasparini et al., 2020b) and it is now established that they play key reproductive roles in all taxa examined (Firman et al., 2017, Fernlund Isaksson and Fitzpatrick, 2023, Hadlow et al., 2023, Zadmajid et al., 2019). This interest is also reflected in recent studies showing that variation in FRF composition is associated to differences in the duration of egg viability (Gueho et al., 2024) and sperm motility parameters (Johnson et al., 2020, Beirão et al., 2015). However, a unified framework linking FRF composition to its action on gametes in general is still lacking, and it currently remains difficult to separate the mechanisms behind FRF influence on eggs from those on sperm.

Regarding sperm attraction specifically, the active compounds identified mostly consist of amino-acids, peptides and proteins (Kholodnyy et al., 2020, Yoshida et al., 2008), whose acquisition and/or synthesis are likely costly to females. The few studies that have directly explored the effects of resource availability on FRF quality have focused on internally fertilising fish and provided mixed results (Cardozo and Pilastro, 2018, Fernlund Isaksson and Fitzpatrick, 2023), but a few indirect lines of evidence suggest that the quality of FRFs, and by extension their functions, may be affected by female condition. For example, ovarian fluid production in medaka is linked to the metabolic activity of follicular cells (Yamamoto, 1963). Similarly, ovarian fluid from farmed Atlantic cod negatively affects the sperm function of wild-caught males (Beirão et al., 2014), most likely due to reproductive dysfunctions in farmed fish. Thus, it makes intuitive sense that an evolutionary loss of the sperm attraction function of FRF, if costly, should be beneficial in bitterling, where gamete encounters are facilitated by mussel respiration.

However, it could also be argued that the water flow inside the mussel may not necessarily translate to a straightforward trajectory of the ejaculate to the site of fertilisation. For instance, female bitterling prefer mussels with a high ventilation rate, which ensures a better oxygen supply to eggs and embryos (Mills and Reynolds, 2002), but could also lead to a shorter duration of sperm retention within the gills. Under these circumstances, active sperm attraction by FRF might be required for gametes to meet within the fertilisation window. In this regard, our preliminary observations suggest that in *R. amarus* sperm longevity is enhanced in a medium containing FRF compared to a water control (Figure 3). This indicates that bitterling FRF may be similar to that of taxa with conventional fertilisation strategies regarding its influence on sperm function.

In summary, we cannot conclude from the current experimental design whether the sperm attraction and longevity enhancement properties of bitterling FRF are required for fertilisation, are maintained by pleiotropic effects of genes governing the primary functions of FRF (support of maturing oocytes) or represent merely an evolutionary artefact. These aspects warrant further investigation in relation to the role of mussel respiration in sperm transport, the pre-oviposition ejaculation behaviour of male bitterling and the naturally long motility duration of bitterling sperm (Smith et al., 2014, Browne et al., 2015).

The sperm attraction property of bitterling FRF revealed here implies that FRF could function as a mediator of cryptic female choice in this species. Male bitterling frequently sneak ejaculations over mussels defended by rivals (Smith and Reichard, 2005). Through preferential boost by FRF of sperm with certain traits (Cattelan et al., 2023) or of certain males (Daupagne et al., 2024, Poli et al., 2019), female bitterling could still bias fertilisation towards preferred or compatible males (Reichard et al., 2012, Smith et al., 2018).

Through their influence on sperm function (Zadmajid et al., 2019), FRFs can affect the outcomes of sperm competition (Gasparini et al., 2020b), which in turn can influence evolutionary processes. For instance, guppy FRF favours sperm of males originating from the same population, indicating that it may function as a post-mating pre-zygotic barrier, ultimately facilitating speciation (Devigili et al., 2018). We found that *R. amarus* sperm is attracted by the FRF of allopatric *S. microlepis* to a higher extent than to a water control (Table 2; Figure 2C-D). Also, *R. amarus* sperm showed no clear preference for the FRF of either species (Table 2; Figure 2E-F). This result is in line with a recent finding from native North American salmon and char and invasive European brown trout, according to which ovarian fluid of native fish improves sperm performance of invasive species to an even higher extent compared to that of conspecifics, despite hybrids being non-viable or sterile (Lantiegne and Purchase, 2023). This could be explained as a side effect of an inbreeding-avoidance strategy of the native species, also seen in a swordtail species complex (E. Morbiato 2026 *In Press*). Alternatively, because the native species have no historical exposure to a closely related heterospecific fish, mechanisms of cryptic female choice might be lagging behind discriminating against heterospecific sperm (Lantiegne and Purchase, 2023). This is in accordance with evidence from fish and bird hybrid zones indicating that the strength of FRF preference for conspecific sperm is proportional to the risk of hybridisation (Yeates et al., 2013, Cramer et al., 2016a, Cramer et al., 2016b). Thus, the most likely explanation for our finding is that mechanisms of heterospecific gamete recognition are not relevant for *R. amarus* because it does not coexist with any other bitterling species.

In conclusion, we found that despite a long evolutionary history of mussel parasitism leading to fertilisation being facilitated by mussel respiration, bitterling FRF actively attracts sperm and considerably extends motility duration. The specific ways in which the mussel microenvironment affects interactions between ejaculates and eggs/FRF remain to be tested.

### Ethics statement

All experiments were performed in accordance with institutional and national guidelines and regulations. Sperm and eggs/fluid extraction were performed by experienced operators according to established protocols. All fish recovered well from anaesthesia and resumed normal swimming within two minutes.

### Data accessibility

The dataset and R script used for generating this paper are available on Figshare (doi: 10.6084/m9.figshare.32790780).

## Acknowledgments

We thank Alessandro Devigili for advice on selection chamber production. We are grateful to Adam Veleba for overseeing the 3D-printing of the sperm selection devices and to Reichard Lab members for helpful discussions. This project was funded by the Czech Science Foundation (grant number 21-00788X to M.R.)

## References

Aguilar, J. & Reyley, M. 2005. The uterine tubal fluid: secretion, composition and biological effects. Animal Reproduction (AR), 2, 91–105.

Alonzo, S. H., Stiver, K. A. & MARSH-Rollo, S. E. 2016. Ovarian fluid allows directional cryptic female choice despite external fertilization. Nat Commun, 7, 12452.

Baer, B., Eubel, H., Taylor, N. L., O’toole, N. & Millar, A. H. 2009. Insights into female sperm storage from the spermathecal fluid proteome of the honeybee Apis mellifera. Genome Biology, 10, R67.

Beirão, J., Purchase, C., Wringe, B. & Fleming, I. 2014. Wild Atlantic cod sperm motility is negatively affected by ovarian fluid of farmed females. Aquaculture Environment Interactions, 5, 61–70.

Beirão, J., Purchase, C. F., Wringe, B. F. & Fleming, I. A. 2015. Inter-population ovarian fluid variation differentially modulates sperm motility in Atlantic cod *Gadus morhua*. Journal of Fish Biology, 87, 54–68.

Browne, R. K., Kaurova, S. A., Uteshev, V. K., Shishova, N. V., Mcginnity, D., Figiel, C. R., Mansour, N., Agnew, D., Wu, M., Gakhova, E. N., Dzyuba, B. & Cosson, J. 2015. Sperm motility of externally fertilizing fish and amphibians. Theriogenology, 83, 1–13.e8.

Candolin, U. & Reynolds, J. D 2002. Adjustments of ejaculation rates in response to risk of sperm competition in a fish, the bitterling (*Rhodeus sericeus*). Proc Biol Sci, 269, 1549–1553.

Cardozo, G. & Pilastro, A. 2018. Female nutritional condition affects ovarian fluid quality in guppies. Biol Lett, 14.

Cattelan, S., Devigili, A., Santacà, M. & Gasparini, C. 2023. Female reproductive fluid attracts more and better sperm: implications for within-ejaculate cryptic female choice. Biology Letters, 19, 20230063.

Chang, C.-H., Li, F., Shao, K.-T., Lin, Y.-S., Morosawa, T., Kim, S., Koo, H., Kim, W., Lee, J.-S., He, S., Smith, C., Reichard, M., Miya, M., Sado, T., Uehara, K., Lavoué, S., Chen, W.-J. & Mayden, R. L. 2014. Phylogenetic relationships of Acheilognathidae (Cypriniformes: Cyprinoidea) as revealed from evidence of both nuclear and mitochondrial gene sequence variation: Evidence for necessary taxonomic revision in the family and the identification of cryptic species. Molecular Phylogenetics and Evolution, 81, 182–194.

Cramer, E. R. A., Ålund, M., Mcfarlane, S. E., Johnsen, A. & Qvarnström, A. 2016a. Females discriminate against heterospecific sperm in a natural hybrid zone. Evolution, 70, 1844–1855.

Cramer, E. R. A., Stensrud, E., Marthinsen, G., Hogner, S., Johannessen, L. E., Laskemoen, T., Eybert, M. C., Slagsvold, T., Lifjeld, J. T. & Johnsen, A. 2016b. Sperm performance in conspecific and heterospecific female fluid. Ecology and Evolution, 6, 1363–1377.

Daupagne, L., Winkler, L., PEMBURY-Smith, M. Q., Lüpold, S., Snook, R. R. & Fitzpatrick, J. L. 2024. One size does not fit all: female–male interactions on the path to fertilization. Reproduction, 169.

Devigili, A., Cattelan, S. & Gasparini, C. 2021. Sperm accumulation induced by the female reproductive fluid: Putative evidence of chemoattraction using a new tool. Cells, 10, 2472.

Devigili, A., Fitzpatrick, J. L., Gasparini, C., Ramnarine, I. W., Pilastro, A. & Evans, J. P. 2018. Possible glimpses into early speciation: the effect of ovarian fluid on sperm velocity accords with post-copulatory isolation between two guppy populations. Journal of Evolutionary Biology, 31, 66–74.

Dietrich, M. A., Dabrowski, K., Arslan, M., Ware, K. & VAN Tassell, J. 2012. Quantifying quality attributes of walleye eggs prior to fertilization—Impact of time of ovulation and gametes storage. Journal of Great Lakes Research, 38, 445–450.

Evans, J. P. & Sherman, C. D. 2013. Sexual selection and the evolution of egg-sperm interactions in broadcast-spawning invertebrates. The Biological Bulletin, 224, 166–183.

FERNLUND Isaksson, E. & Fitzpatrick, J. L. 2023. Examining the potential for resource-dependent female reproductive fluid-sperm interactive effects in a livebearing fish. Journal of Evolutionary Biology, 36, 709–719.

Firman, R. C., Gasparini, C., Manier, M. K. & Pizzari, T. 2017. Postmating Female Control: 20 Years of Cryptic Female Choice. Trends Ecol Evol, 32, 368–382.

Fitzpatrick, J. L., Willis, C., Devigili, A., Young, A., Carroll, M., Hunter, H. R. & Brison, D. R. 2020. Chemical signals from eggs facilitate cryptic female choice in humans. Proceedings of the Royal Society B: Biological Sciences, 287, 20200805.

Gasparini, C., Pilastro, A. & Evans, J. P. 2020a. The role of female reproductive fluid in sperm competition. Philosophical Transactions of the Royal Society B-Biological Sciences, 375.

Gasparini, C., Pilastro, A. & Evans, J. P. 2020b. The role of female reproductive fluid in sperm competition. Philosophical Transactions of the Royal Society B: Biological Sciences, 375, 20200077.

Gueho, A., Żarski, D., Rime, H., Guével, B., Com, E., Lavigne, R., Nguyen, T., Montfort, J., Pineau, C. & Bobe, J. 2024. Evolutionarily conserved ovarian fluid proteins are responsible for extending egg viability in salmonid fish. Scientific Reports, 14.

Hadlow, J. H., Evans, J. P. & Lymbery, R. A. 2023. Female reproductive fluids ‘rescue’ sperm from phenotypic ageing in an external fertilizer. Proceedings of the Royal Society B: Biological Sciences, 290, 20230574.

Hartig, F. 2016. DHARMa: residual diagnostics for hierarchical (multi-level/mixed) regression models. CRAN: Contributed Packages.

Heschl, A. 1989. Integration of “innate” and “learned” components within the IRME for mussel recognition in the European bitterling Rhodeus amarus (Bloch). Ethology, 81, 193–208.

Johnson, S. L., Borziak, K., Kleffmann, T., Rosengrave, P., Dorus, S. & Gemmell, N. J. 2020. Ovarian fluid proteome variation associates with sperm swimming speed in an externally fertilizing fish. Journal of Evolutionary Biology.

Johnson, S. L., Villarroel, M., Rosengrave, P., Carne, A., Kleffmann, T., Lokman, P. M. & Gemmell, N. J. 2014. Proteomic Analysis of Chinook Salmon (Oncorhynchus tshawytscha) Ovarian Fluid. PLoS One, 9, e104155.

Kholodnyy, V., Gadêlha, H., Cosson, J. & Boryshpolets, S. 2020. How do freshwater fish sperm find the egg? The physicochemical factors guiding the gamete encounters of externally fertilizing freshwater fish. Reviews in Aquaculture, 12, 1165–1192.

Kitamura, J.-I. 2007. Reproductive Ecology and Host Utilization of Four Sympatric Bitterling (Acheilognathinae, Cyprinidae) in a Lowland Reach of the Harai River in Mie, Japan. Environmental Biology of Fishes, 78, 37–55.

Kustra, M. C., Servedio, M. R. & Alonzo, S. H. 2025. Cryptic female choice can maintain reproductive isolation. Evolution, 79, 2259–2273.

Lantiegne, T. H. & Purchase, C. F. 2023. Can cryptic female choice prevent invasive hybridization in external fertilizing fish? Evolutionary Applications, 16, 1412–1421.

Lüdecke, D., BEN-Shachar, M., Patil, I., Waggoner, P. & Makowski, D. 2021. performance: An R Package for Assessment, Comparison and Testing of Statistical Models. Journal of Open Source Software, 6, 3139.

Lymbery, R. A., Kennington, W. J. & Evans, J. P. 2017. Egg chemoattractants moderate intraspecific sperm competition. Evolution Letters, 1, 317–327.

Mcgillycuddy, M., Warton, D. I., Popovic, G. & Bolker, B. M. 2025. Parsimoniously Fitting Large Multivariate Random Effects in glmmTMB. Journal of Statistical Software, 112.

Mills, S. C. & Reynolds, J. D. 2002. Mussel ventilation rates as a proximate cue for host selection by bitterling, Rhodeus sericeus. Oecologia, 131, 473–478.

Myers, J. N., Bradford, A. J., Hallas, V. S., Lawson, L. L., Pitcher, T. E., Dunham, R. A. & Butts, I. A. E. 2020. Channel catfish ovarian fluid differentially enhances blue catfish sperm performance. Theriogenology, 149, 62–71.

Nakagawa, S., Schielzeth, H. & O’hara, R. B. 2013. A general and simple method for obtainingR2from generalized linear mixed-effects models. Methods in Ecology and Evolution, 4, 133–142.

Pinzoni, L., Poli, F., Grapputo, A., Rasotto, M. B. & Gasparini, C. 2023. Female Reproductive Fluid Increases the Opportunities for Postmating Sexual Selection by Prolonging the Egg Fertilization Window. The American Naturalist, 201, 491–499.

Pinzoni, L., Rasotto, M. B. & Gasparini, C. 2024. Sperm performance in the race for fertilization, the influence of female reproductive fluid. Royal Society Open Science, 11, 240156.

Poli, F., Immler, S. & Gasparini, C. 2019. Effects of ovarian fluid on sperm traits and its implications for cryptic female choice in zebrafish. Behavioral Ecology, 30, 1298–1305.

Reichard, M., Smith, C. & Jordan, W. C. 2004. Genetic evidence reveals density-dependent mediated success of alternative mating behaviours in the European bitterling *Rhodeus sericeus*. Molecular Ecology, 13, 1569–1578.

Reichard, M., Spence, R., Bryjová, A., Bryja, J. & Smith, C. 2012. Female Rose Bitterling Prefer MHC-Dissimilar Males: Experimental Evidence. PLoS One, 7, e40780.

Serrano, H., Canchola, E. & GARCÍA-Suárez, M. D. 2001. Sperm-Attracting Activity in Follicular Fluid Associated to an 8.6-kDa Protein. Biochemical and Biophysical Research Communications, 283, 782–784.

Smith, C. & Reichard, M. 2005. Females solicit sneakers to improve fertilization success in the bitterling fish (Rhodeus sericeus). Proceedings of the Royal Society B: Biological Sciences, 272, 1683–1688.

Smith, C., Reichard, M. & Jurajda, P. 2003. Assessment of sperm competition by European bitterling, Rhodeus sericeus. Behavioral Ecology and Sociobiology, 53, 206–213.

Smith, C., Reichard, M., Jurajda, P. & Przybylski, M. 2004. The reproductive ecology of the European bitterling (Rhodeus sericeus). Journal of Zoology, 262, 107–124.

Smith, C., Spence, R. & Reichard, M. 2018. Sperm is a sexual ornament in rose bitterling. Journal of Evolutionary Biology, 31, 1610–1622.

Smith, C., Warren, M., Rouchet, R. & Reichard, M. 2014. The function of multiple ejaculations in bitterling. Journal of Evolutionary Biology, 27, 1819–1829.

Yamamoto, K. 1963. Cyclical Changes in the Wall of the Ovarian Lumen in the Medaka, Oryzias latipes. Annotationes zoologicae Japonenses, 36, 179–186.

Yeates, S. E., Diamond, S. E., Einum, S., Emerson, B. C., Holt, W. V. & Gage, M. J. G. 2013. CRYPTIC CHOICE OF CONSPECIFIC SPERM CONTROLLED BY THE IMPACT OF OVARIAN FLUID ON SPERM SWIMMING BEHAVIOR. Evolution, 67, 3523–3536.

Yoshida, M., Kawano, N. & Yoshida, K. 2008. Control of sperm motility and fertility: Diverse factors and common mechanisms. Cellular and Molecular Life Sciences, 65, 3446–3457.

Yoshida, M., Murata, M., Inaba, K. & Morisawa, M. 2002. A chemoattractant for ascidian spermatozoa is a sulfated steroid. Proceedings of the National Academy of Sciences, 99, 14831–14836.

Zadmajid, V., Myers, J. N., Sørensen, S. R. & Butts, I. A. E. 2019. Ovarian fluid and its impacts on spermatozoa performance in fish: A review. Theriogenology, 132, 144–152.

Zhao, J., Xu, D., Zhao, K., Diogo, R., Yang, J. & Peng, Z. 2016. The origin and divergence of Gobioninae fishes (Teleostei: Cyprinidae) based on complete mitochondrial genome sequences. Journal of Applied Ichthyology, 32, 32–39.

